# Iron acquisition across outer membrane: The role of major outer membrane protein Slr1908 under iron supplemented conditions in *Synechocystis* 6803

**DOI:** 10.1101/2021.01.14.426630

**Authors:** Rachna Agarwal, Sharmistha Dutta Choudhury, Akipeddi V S S Narayana Rao

## Abstract

The Ton-B dependent outer membrane (OM) transporters are responsible for active iron (Fe) import in *Synechocystis* sp. strain PCC 6803 (*S*. 6803 or WT) under Fe depletion. However, the mechanism of Fe acquisition under Fe supplemented conditions remains uncharacterised. In the present study, functional role of OMP Slr1908 in *S*. 6803 was addressed by insertional mutagenesis. The *Δslr1908* cells exhibited slower growth in the first week in comparison to the WT and displayed an absorption and 77K fluorescence spectrum typical of Fe deficiency. Indeed, the mutant had ∼ 80% less Fe as confirmed by atomic absorption spectroscopy and ^55^Fe-radiotracer uptake. The iron deficiency was paralleled with low Mn content. The mutant had low SOD content as well as activity, less cytochromes, less chlorophyll content, less F_v_/ F_m_, lower ETRII and high oxidative stress in comparison to the WT at the end of first week. Interestingly, the mutant showed transcriptional upregulation of iron stress induced protein *isiA* and *isiB* signifying intracellular Fe deficiency. Upregulation of OMP Slr0042 was also observed at RNA and protein level. The results indicate that Slr1908 is a major Fe uptake OMP in *S*. 6803 the deletion of which leads to initial slow growth that gets partially offset by induction of other Fe importing OMPs.

## 1. Introduction

Iron (Fe) is a key inorganic micronutrient for all living organisms [1-2]. It is an essential co-factor for redox centres of oxidative phosphorylation, chlorophyll biosynthesis, oxidative stress detoxification and so on [1-2]. Iron is the fourth most abundant element in the earth’s crust. However, it has poor bioavailability as it gets readily oxidised to water insoluble ferric hydroxide especially in the aquatic systems due to the oxidising nature of the earth’s environment [3]. Hence, living organisms have devised specialised mechanisms such as siderophore secretion for efficient Fe intake for optimum metabolism [4]. They have also evolved dedicated adaptation strategies to cope up with Fe limitation and survive through the unfavourable conditions [3, 5].

Fe requirement by photoautotrophs such as cyanobacteria is typically ten times higher than heterotrophs owing to a large amounts required for photosynthetic light harvesting [6]. Cyanobacteria account for 25% of global primary productivity and require large Fe pools [7]. Like Gram-negative bacteria, they have a multi-layered cell envelop composed of a plasma membrane, a peptidoglycan cell wall and an outer membrane (OM) [8]. OMs are asymmetrical lipid bilayers, having trans-membrane outer membrane proteins (OMPs) with exclusive β-barrel secondary structure which are the first port of entry for all the inorganic nutrients required for metabolism [9]. *S*. 6803 does not synthesize and secrete siderophore for Fe uptake and hence primarily utilises the inorganic Fe present in the environment [10-11]. Fe transport through bacterial inner membrane is facilitated through multiple pathways such as Fut, Feo and ExbB-ExbD (OM energisation system) [10, 12]. In the case of *S*. 6803, OM localised TonB dependent transporters (TBDTs) works in conjunction with ExbB-ExbD and are important under iron depleted environmental conditions [13-14]. The other route for Fe import in this organism may be OM porins namely *sll0772, sll1271, sll1550, slr0042, slr1272, slr1841 and slr1908* (Table S1). Out of these, four minor OMPs (Sll0772, Sll1271, Sll1550 and Slr0042) are required for Fe uptake under Fe deficiency [14]. However, there are no reported dedicated routes for routine Fe uptake under well supplemented growth conditions [10].

*slr1908* that encodes a putative OMP is a highly expressed gene and the corresponding protein has been shown to be present exclusively in the purified OM of *S*. 6803 [15-16]. Functionally, Slr1908 protein has been shown to be impermeable to organic nutrients and allows the passage of only small inorganic ions in vitro [17]. Previous report of generating a combined homozygous OMP mutant involving *slr1908* have been unsuccessful indicating that the encoded protein might be physiologically indispensable [14]. Interestingly, in the present study, we could obtain a homozygous *Δslr1908* mutant of *S*. 6803 and performed its physiological characterisation. The *Δslr1908* cells grew slowly under Fe supplemented (complete BG-11) medium containing ∼ 23 µM Fe, supplied as ferric ammonium citrate. The mutant also showed characteristic features of Fe deficiency such as induction of *isiA* and *isiB* genes as well as low Fe content and high oxidative stress in spite of being grown in complete BG-11 medium. The results clearly attribute the function of Fe import to Slr1908.

## 2. Materials and methods

### 2.1 Growth media and culture conditions

*S*. 6803 GT (glucose tolerant) strain was grown in BG-11 medium (pH 7.8) or on BG-11 agar plates under continuous white light of intensity ∼ 30 µmoles m^-2^ s^-1^ and temperature between 28-30 °C with continuous shaking (140 rpm) [18]. The Δ*slr1908* mutant strain was grown in BG-11 medium supplemented with 25-100 μg/ ml spectinomycin. *E. coli* DH5α strain was grown on LB plates or broth at 37 °C (under continuous shaking at 180 rpm) supplemented with appropriate antibiotic.

### 2.2 Construction of Δ*slr1908* mutant of *S*. 6803

The Δ*slr1908* mutant of *S*. 6803 was generated according to Agarwal et al. [19] and is described in Fig. 1A. Briefly, the 1776 bp long *slr1908* fragment was amplified using the primers P1 and P2 listed in Table S2 and cloned between NdeI and XhoI sites of plasmid pET23b+ to obtain the plasmid pET-1908. pET-1908 was linearised by double digestion with EcoRV and HpaI releasing a fragment of ∼1079 bp. A 1963 bp blunt-ended spectinomycin resistance cassette was cloned in the linearised pET-1908 to generate the plasmid pET-1908::aad [19]. The size of the final mutated *slr1908* in pET-1908::aad is ∼ 2679 bp. All the cloning procedures were performed in *E. coli* DH5α cells and the recombinants were confirmed by colony PCR. The recombinant plasmid pET-1908::aad was used for transforming *S*. 6803 cells to obtain Δ*slr1908* cells using standard procedures (Fig. 1A) [19]. The selected transformants were segregated to homozygosity by restreaking on the plates containing an increasing concentration of the same antibiotic (10-100 µg/ ml). The segregation status was assessed and confirmed by colony PCR using the gene specific as well as flanking primers listed in Table S2.

**Fig. 1:**
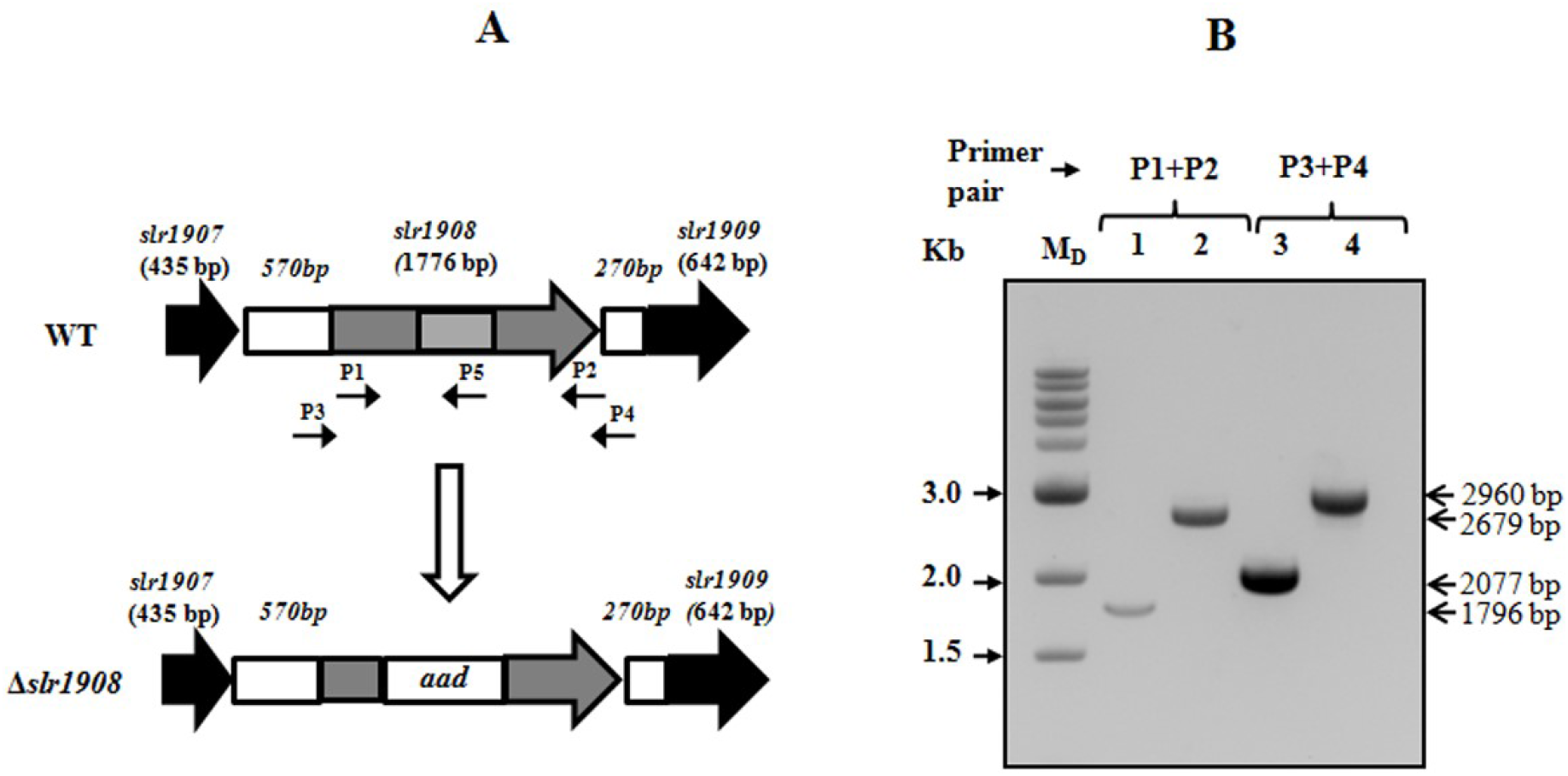
**(A)** Schematic representation of the organisation of *slr1908* locus in WT and *Δslr1908* in *S*. 6803. Figure not to scale. Light gray box denotes the deleted region in WT for generating the *Δslr1908* mutant. **(B)** Confirmation of homozygosity of *Δslr1908* cells by colony PCR and resolving the products on 1% AGE. The primers used for amplification are denoted above the lanes (Table S2). Lanes 1 & 3: WT; Lanes 2 & 4: *Δslr1908*. Size of the bands is indicated on RHS with black arrows. M_D_: DNA ladder

### 2.3 Growth measurement

Wild-type and Δ*slr1908 S*. 6803 cells were inoculated in BG-11 medium with or without antibiotic (Spectinomycin, 25 µg/ ml), with an initial O.D._730 nm_ of 0.05 and were grown as in 2.1. Aliquots were withdrawn at regular intervals to measure O.D._730 nm_ as an indicator of growth.

### 2.4 Absorbance and 77 K fluorescence measurements

Whole cell absorption spectra of wild-type and Δ*slr1908 S*. 6803 cells (O.D. _730 nm_ ∼ 0.6) were recorded between 400-750 nm at room temperature [20] and normalised with respect to the absorbance at 730 nm.

For 77K fluorescence measurement, 1 ml of WT and Δ*slr1908 S*. 6803 cells (O.D._730nm_ of ∼ 0.6) was put in a nuclear magnetic resonance tube (5 mm diameter) and frozen under liquid nitrogen. The emission spectra were recorded under liquid nitrogen at excitation wavelength of 435 nm and emission range of 520-800 nm (excitation and emission slit width of 5 nm each) on a Hitachi spectrofluorimeter (F-4510). The spectra were appropriately normalised in OriginPro Ver. 8.

### 2.5 Chlorophyll determination and pulse amplitude modulated (PAM) fluorimetry

Chlorophyll was determined as described in Agarwal et al [20]. The chlorophyll a content was calculated using the equation:

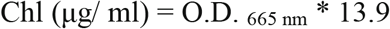

For PAM fluorimetry, cells equivalent to O.D. _730 nm_ of ∼ 2.0 were collected and resuspended in 2 ml fresh BG-11 medium. After 30 min of dark adaptation, the cell suspension was kept in a fluorimetric cuvette (path length 1 cm) on the Dual-PAM instruments’s cuvette holder and the F_v_/ F_m_ as well as rapid light curves were recorded using DUAL-DR emitter detector under stirring.

### 2.6 SDS-PAGE and western blotting

For resolving the total cellular proteins, cells equivalent to an O.D. _730 nm_ ∼25.0 were collected by centrifugation at 10, 000 x g for 2 min at RT, washed once with fresh BG-11 medium, resuspended in ∼ 200 µl of lysis buffer [20 mM Tris-Cl pH 8.0, 50 mM NaCl, 1 mg/ ml lysozyme, 1 mg/ ml DNaseI, 1 mM MgCl_2_, 10 % sucrose, protease inhibitor cocktail (Roche)] and incubated at RT for 1 h. The suspension was mixed with equal volume of 100 µm glass beads and vortexed thrice for 1 min each. The samples were freeze thawed thrice in liquid nitrogen and incubated for 1 h at 4 °C with 1% β-dodecyl maltoside to extract membrane proteins. The final cell lysate was centrifuged at 10, 000 x g for 30 min. The supernatant representing the soluble and extracted membrane proteins was resolved either on 10% SDS-PAGE at RT or 12 % native-PAGE at 4 °C as described previously [21].

For western blotting, 10 µl of the cell lysate was resolved on 12% SDS-PAGE, electro-blotted onto a nitrocellulose membrane, blocked, treated with appropriately diluted primary and secondary antibodies and colorimetrically detected using NBT-BCIP method following manufacturer’s instructions. All primary antibodies were commercially raised in rabbit and the corresponding antigen was over-expressed in *E. coli*. The Mn-SOD antibody was raised against the protein from *Anabaena* PCC 7120 and it showed > 50% identity with the single Fe-SOD from *S*. 6803 [22].

### 2.7 Activity staining for superoxide dismutase and catalase

The whole cell extract was resolved on 10% native-PAGE at 4 °C. For in-gel activity staining of catalase, the gel was incubated with 50 ml of 0.003% H_2_O_2_ for 40 min in dark and developed with a solution containing 1% FeCl_3_ and K_3_Fe(CN)_6_. For in-gel activity staining of superoxide dismutase, the gel was incubated with a solution containing 0.25 mM NBT, 10 mM riboflavin and 28 mM TEMED in dark for 1 h followed by exposing the gel to bright light until appearance of clearance bands against a purple background.

### 2.8 In-gel cytochrome staining and DCHFDA assay for reactive oxygen species

10 µl of the whole cell lysate was resolved on 12% non-reducing SDS-PAGE and the staining was done according to Hansen (1980) [23].

DCHFDA assay was performed essentially according to Agarwal et al (2008) and the fluorescence was measured in a BioTek multimode plate reader [24].

### 2.9 Immunofluorescence

For immuno-detection of Slr1908 protein, anti-Slr1908 and anti-Sll1951 antibody was commercially generated in rabbit (Bangalore Genie Pvt. Ltd., India). The immunofluorescence experiment was performed according to Rast et al. [25]. The overnight cured slides were viewed under an Olympus IX83 research inverted fluorescence microscope using appropriate filters. The images were analyzed using the Adobe Photoshop 7.0 program. ∼600 cells were viewed for the WT and the mutants in random fields.

### 2.10 Transmission Electron Microscopy (TEM)

The samples for TEM were processed according to Agarwal et al [19]. Final embedding and polymerization were performed in Spurr’s resin at 72 °C for 48 h. The 70 nm ultrathin sections were taken on 100-mesh copper grids, stained with 2% uranyl acetate (aq.) for 30 min and viewed under the Zeiss FE-TEM at 200 KeV.

### 2.11 Total RNA isolation and RT-qPCR

Total RNA was isolated from ∼ 45 ml of one week old cultures of wild-type and Δ*slr1908 S*. 6803 cells using TRIzol™ LS reagent (ThermoFisher Scientific) following manufacturer’s instructions. Briefly, the cells were pelleted by centrifugation at 5, 000 x g at 24 °C and resuspended in 250 μl of BG-11 medium followed by addition of 750 μl of TRIzol™ LS reagent and 100 μm glass beads. The cells were broken by bead beating thrice at 6 m/s with intermittent cooling on ice. Total RNA was subsequently isolated following chloroform extraction, washed with 70% ethanol and dissolved in 50 μl water. RNA quality was assessed by measuring O.D. 260/ 280 and O.D. 260/ 230 as well as resolving 1 μg of total RNA on a non-denaturing TBE gel. DNase I treatment was given to avoid genomic DNA contamination followed by purification of total RNA using GenElute™ Total RNA Purification Kit (Sigma-Aldrich). The purified RNA was quantified using a NanoDrop 2000 spectrophotometer (ThermoFisher Scientific) and ∼ 90 ng was used for one-step RT-qPCR using Luna® Universal one-step RT-qPCR kit (New England BioLabs Inc.) [10 min. RT step at 55 °C, denaturation at 95 °C for 1 min followed by 45 cycles of three step PCR (Denaturation at 95 °C for 15 s, annealing at 55 °C for 15 s, extension at 72 °C for 30 s) and a high resolution melt curve (HRM) in the end] using the primers listed in Table S2 on a Rotor-Gene Q instrument (Qiagen). *rnpB* was taken as reference for all genes. Primer specificity was ascertained by PCR prior to RT-qPCR using genomic DNA as template (Fig S2). Relative transcript quantitation was performed by 2^-ΔΔCt^ method [26]. Statistically significant differences in transcript abundance were arrived at by performing Student’s t (p≤0.05) using OriginPro 8.0.

### 2.12 Atomic Absorption Spectroscopy

For assaying the uptake of six essential trace metals (Ca, Cu, Fe, Mg, Mn, and Zn) by one week and three weeks old cultures of wild-type and Δ*slr1908 S*. 6803 cells, 45 ml culture was pelleted by centrifugation at 5, 000 x g at 24 °C and washed twice with 1 mM EDTA to remove surface bound metal ions. The final wet pellet was air dried, digested with HNO_3_ (AR) and re-dried completely. The digested dried cell material was resuspended in 10 ml 3% HNO_3_ (AR) and analysed on AAS. All samples were analysed in biological and technical triplicates and the results were expressed as ppm/ mg dry weight.

### 2.13 Iron uptake assays using ^55^FeCl_3_

One week old cultures of WT and Δ*slr1908* cells were washed twice with 5 ml BG-11 medium devoid of citrate and Fe (BG-11_CF_) and resuspended in the same medium to a final O.D._730 nm_ of 1. Fe was supplied as ^55^FeCl_3_ (specific activity 37.2 mCi/ g Fe, obtained from Board of Radiation and Isotope Technology, Mumbai). Sodium ascorbate and ferrozine were added for keeping Fe in ferrous (Fe^2+^) and ferric (Fe^3+^) state respectively (molar ratio of Fe to ascorbate was 1:10 and Fe to ferrozine was 1:2). For analysing the Fe uptake in complex form, desferoxamine B was used (molar ratio of Fe to DFB was 1:2). The uptake was followed for 30 min. in light and the reaction was quenched by adding 10 µl of 0.5 M EDTA (pH 8.0), filtered through 0.45 µm filter discs and immediately washing the filter discs with ice cold BG-11_CF_. The cells on the filter discs were digested with HNO_3_ (AR) in glass scintillation vials and evaporated to dryness. The ^55^Fe uptake was measured as counts per minute in a liquid scintillation counter with 0.3% BBOT in toluene as the scintillation fluor. All samples were analysed in biological as well as technical triplicates and the results were expressed as counts/ O.D._730 nm_/ h.

## 3. Results

### 3.1 Confirmation of homozygous Δ*slr1908* mutant

The segregation status of Δ*slr1908* cells was confirmed by colony PCR with two different primer combinations P1 and P2 (gene specific strict primers at the beginning and end of the ORF respectively), P3 and P4 (external primers) (Table S2, Fig. 1). The WT cells gave single band at ∼1796 bp (primers P1 and P2) and 2077 bp (primers P3 and P4) whereas Δ*slr1908* cells showed single bands at 2679 bp (primers P1 and P2) and at 2960 bp (primers P3 and P4) (Fig. 1B). Further, RT-PCR with a 20 bases long reverse primer positioned in the deleted region of the mutant (P5) (Fig. 1A) also confirmed complete segregation in the latter. The results showed a band of ∼ 564 bp (primers P1 and P5) and a band of ∼684 bp (primers P3 and P5) in RT-PCR of WT cells only (Fig. 2A, Lanes 1 and 3). No corresponding or any other band was observed in Δ*slr1908* cells (Fig. 2A, Lanes 2 and 4). As the region complementary to the reverse primer (P5) has been deleted in the Δ*slr1908* cells, it is expected that a completely homozygous mutant will not be able to synthesise any cDNA and hence no PCR amplification should be observed (Fig. 2A, Lanes 2 and 4). *rnpB* was taken as the housekeeping gene that was expressed in both WT and Δ*slr1908* cells in equivalent amount on equal RNA basis (Fig. 2A, Lanes 5 and 6).

**Fig. 2:**
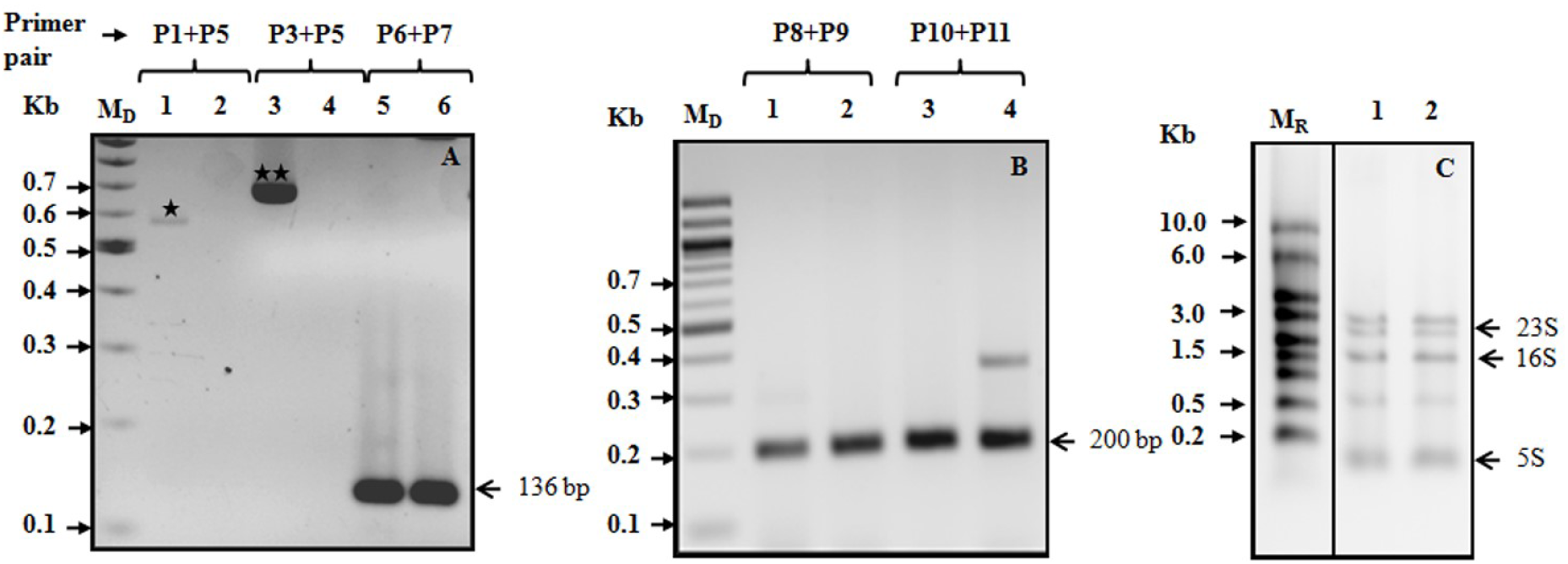
**(A)** RT-PCR of WT and *Δslr1908* cells. cDNA was synthesized from total RNA using a 20 bp reverse primer (P5) complementary to the deleted region in *Δslr1908* cells followed by PCR with primers as indicated above the lanes. The amplified PCR product was resolved on 2% AGE. Lanes 1 & 2: WT and *Δslr1908* cells with primers P1 and P5 respectively (indicated by single black star); Lanes 3 & 4: WT and *Δslr1908* cells with primers P3 and P5 respectively (indicated by two black stars); Lanes 5 & 6: WT and *Δslr1908* cells with primers P6 & P7 respectively (for *rnpB* reference gene). **(B)** Assessment of transcription status of the adjacent genes *slr1907* and *slr1909* in the *Δslr1908* cells by RT-qPCR (Ct values have been mentioned in the text). The PCR products were resolved on 2% AGE. Lanes 1 & 2: WT and *Δslr1908* cells with primers P8 and P9 respectively (for *slr1907*); Lanes 3 & 4: WT and *Δslr1908* cells with primers P10 and P11 (for *slr1909*). The band position is indicated on RHS with a black arrow. **(C)** 1 µg of total RNA was resolved on 1% TBE gel for checking RNA integrity and equivalence. Lane 1: WT; Lane 2: *Δslr1908*. Bands corresponding to rRNA are indicated on RHS with black arrow. Primers P1-P11 are defined with sequence in table S2. Kb: kilobase; bp: base pair: M_D_: DNA ladder; M_R_: RNA ladder

Prokaryotic genes are often transcribed as polycistronic mRNA encoding multiple proteins (operons). Kopf et al (2014) had shown that *slr1908* is transcribed as a single transcriptional unit (with a 120 bp 5’ UTR) along with *slr1909* and *slr1910* (TU599 with 610937 as the transcription start site and 614120 as the end site) [27]. Hence, interruption of *slr1908* may lead to polar effects on the expression of the downstream genes *slr1909* and *slr1910* as well as the upstream gene *slr1907*. RT-qPCR analysis of these two loci i.e. *slr1907* and *slr1909* in WT and Δ*slr1908* cells showed that the two genes are expressed in equivalent amounts in both the cell types on equal RNA basis (Ct values for *slr1907* were 25.9 and 26.8 whereas for *slr1909* they were 25.4 and 26.3 for the WT and the Δ*slr1908* cells) (Fig. 2B lane 1-4, Fig. 2C). It should be appreciated that there exists an alternate transcription start site in TU599 immediately after *slr1908* at position 613142 that can lead to uninterrupted transcription of *slr1909* (and may be *slr1910*) in spite of interruption in *slr1908* [27]. Hence, the phenotypic effects discussed later in this article can be specifically ascribed to absence of functional Slr1908 protein only.

### 3.2 The Δ*slr1908* cells are viable but slow growing and have less chlorophyll

The Δ*slr1908* cells looked pale with as well as without antibiotic in the initial phase (15 days in case of solid plates, 1 week in case of liquid medium) indicating altered pigment composition of the mutant (Fig. 3A panel 1 and 3, Fig. 3B). Nevertheless, longer incubation of 55 days on plates and ∼ 3 weeks in liquid medium showed no visible difference in the colour of the culture (Fig. 3A panel 3 and 4, Fig. 3B). Further, the mutant when grown in presence or absence of the antibiotic showed a similar visible growth pattern indicating that the antibiotic doesn’t have a role in imparting the slow growing pale phenotype to the mutant (Fig. 3B). The growth curve also corroborated that the Δ*slr1908* cells grew slowly in comparison to the WT cells in the beginning (Fig. 3C). The WT achieved an O.D. _730 nm_ of ∼ 0.9-1.0 whereas the Δ*slr1908* had O.D. _730 nm_ of ∼ 0.25-0.3 at the end of the first week when both were inoculated with an initial O.D. _730 nm_ of 0.05. However, Δ*slr1908* mutant resumed faster growth at the later stage (between day 7 and day 17 i.e. second and third week), indicating that the cells were readjusting their physiology in response to the loss of functional Slr1908 (Fig. 3C). Chlorophyll determination on equal cell density basis for one week old cultures showed ∼3.8 µg Chl/ O.D. _730nm_ for the WT and ∼2.6-2.7 µg Chl/ O.D. _730 nm_ for the Δ*slr1908* cells (∼31% less chlorophyll in the mutant) that correlated with the mutant’s pale phenotype, slow initial growth and further corroborated its altered physiology (Fig. 3D).

**Fig. 3:**
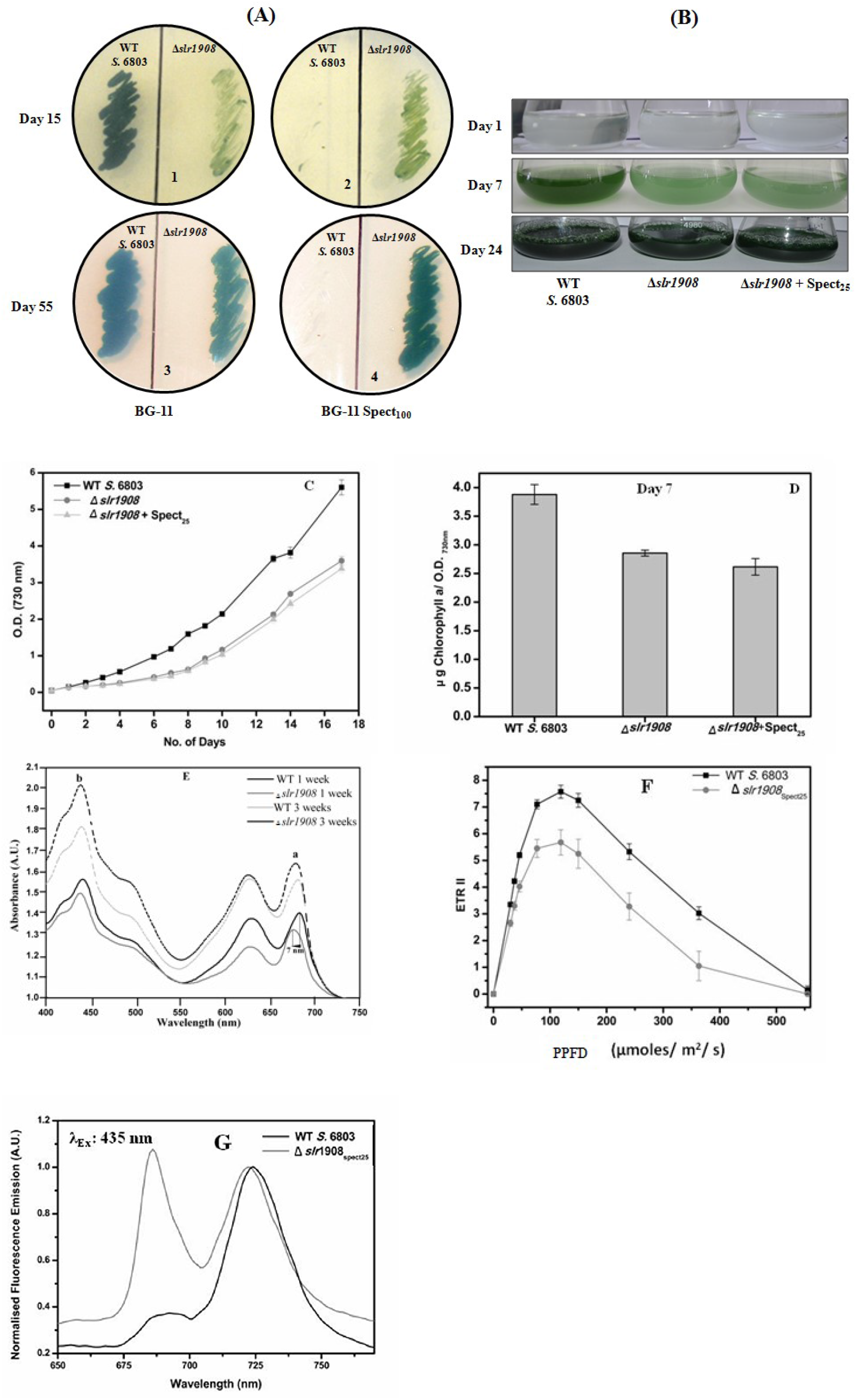
**(A)** Growth of WT and *Δslr1908* cells on BG-11 as well as BG-11-_Spect100_ plates **(B)** Growth in BG-11 medium with and without spectinomycin (25 µg/ ml). **(D)** Growth curve of WT and *Δslr1908* cells. **(D)** Chlorophyll content of WT and *Δslr1908* cells after 7 days of growth in liquid BG-11 medium. **(E)** Visible light absorption spectrum of WT and *Δslr1908* cells after one week and three weeks of growth. Chlorophyll absorption peak in the red region (a) and blue region (b) of the visible spectrum are marked. The 7 nm blue shift in chlorophyll absorption in red region in *Δslr1908* cells with respect to WT cells is marked by a black arrow. **(F)** Rapid light curve of one week old WT and *Δslr1908* cells showing electron transport through photosystem II (ETRII) **(F)** Low temperature (77K) fluorescence emission spectrum of WT and *Δslr1908* cells at the end of first week of growth (excitation at 435 nm). Spectra have been normalised with respect to PSI emission peak (720-725 nm). All experiments were repeated three or more times independently and representative data has been shown. Values represent Average±SEM (n=3), wherever applicable.

### 3.3 The Δ*slr1908* cells showed a blue shift in the chlorophyll absorption spectrum, poor F_v_/F_m_ and ETRII as well as higher PSII fluorescence at 77K

The absorption spectrum recorded between 400-750 nm showed a decreased peak height for chlorophyll absorption and a prominent ∼ 7 nm blue shift in the chlorophyll absorption peak in the red region of the spectrum in the Δ*slr1908* cells with respect to the WT (Fig. 3E). While WT showed a chlorophyll absorption maximum at ∼ 441 nm and ∼ 682 nm respectively (both after 1 week and 3 weeks), the mutant had peaks at 439 nm and 675 nm (7 nm shift) after one week and 439 nm and 678 nm (3 nm shift) after 3 weeks. The blue shift in the chlorophyll absorption peak has been ascribed to iron deficiency in many cyanobacteria. In the present case, an initial (after first week of inoculation) prominent blue shift in the red region of the spectrum that became less pronounced as the growth continued may imply a gradual physiological adaptation of the mutant as the growth continues (discussed later). At the end of first week of growth, the Δ*slr1908* cells showed ∼ 11% lower maximum quantum efficiency of PSII (F_v_/F_m_) (0.301±0.008) in comparison to the WT (0.337±0.029) whereas they had comparable values after three weeks of growth (0.299±0.008 for WT and 0.310±0.004 for the mutant). Further, the rapid light curves indicated similar rise in ETR(II) with saturation of electron transport rate at a PPFD of ∼ 100 µmoles/ m^2^ / s and subsequent photo-inhibition occurring beyond PPFD of ∼ 150 µmoles/ m^2^ / s for WT and Δ*slr1908* cells (Fig. 3F). However, the maximum ETRII of the Δ*slr1908* cells was ∼ 25% lower in comparison to the WT indicating its poor photosynthetic performance (Fig. 3F). Low temperature (77K) fluorescence emission spectrum obtained after exciting chlorophylls at 435 nm (normalised to emission at peak corresponding to PSI at 721-725 nm) showed that mutant cells had unusually high emission from PSII at 685 nm and 695 nm respectively in comparison to the WT (Fig. 3G). Further, the chlorophyll emission was blue shifted in case of Δ*slr1908* cells (∼721 nm) in comparison to the WT (∼ 725 nm) (Fig. 3G). These are the hallmark features reported in the case of iron deficiency in many cyanobacteria [28-29]. The results imply that the relatively poor photosynthetic performance of the *Δslr1908* cells might be due to its poor iron uptake capacity which is an essential micronutrient and an integral component of redox centres of photosynthesis.

### 3.4 Slr1908 loss leads to severely altered ultrastructure in the Δ*slr1908 cells*

Exact cellular location of Slr1908 protein (in the WT) and the deletion confirmation (in Δ*slr1908* cells) was delineated by immunofluorescence using an anti-Slr1908 antibody (Fig. 4A). Anti-Sll1951 antibody labelled cells served as positive control for both cell types (Fig. 4B). Differential interference contrast (DIC) [Fig. 4 A (a and f), 4 B (k and p)], Phycobilisome/Chlorophyll fluorescence [Fig. 4 A (b and g), 4 B (l and q)] and DAPI fluorescence from DNA [Fig. 4 A (c and h), 4 B (m and r)] imaging was employed to locate the cells. Antigen location (Slr1908 or Sll1951) was visualised using Alexa-488 fluorescence [Fig. 4 A (d and i), 4 B (n and s)] followed by merging the three fluorescence channels (Phycobilisome/Chlorophyll, DAPI, Alexa 488) to obtain a consolidated antigen distribution landscape [Fig. 4 A (e and j), 4 B (o and t)]. The results showed 1.5-2 µm sized ellipsoidal/ spherical dividing cells (Fig. 4 A and B). Slr1908 protein could be localised in the cell envelop in discrete regions in WT cells [Fig. 4 A (d and e)] as against Sll1951 that formed a continuos ring in WT as well as Δ*slr1908* cells [Fig. 4 B (n, o, s and t)]. Surprisingly, some unexpected fluorescence was observed in Δ*slr1908* cells as well that have previously been shown to be completely segregated homozygous mutant for *slr1908* [Fig. 4 A (i and j)]. However, *S*. 6803 contains 7 putative OMPs out of which the four proteins Slr1841, Slr1908, Slr0042 and Sll1550 share ∼ 90% identity in the first 120 amino acids at N-terminus (after cleavage of signal peptide) (Fig. S1 A and B). As the primary antibody against Slr1908 was raised using an intact Slr1908 protein, the cross-reactivity with these other OMPs gets substantiated.

**Fig. 4:**
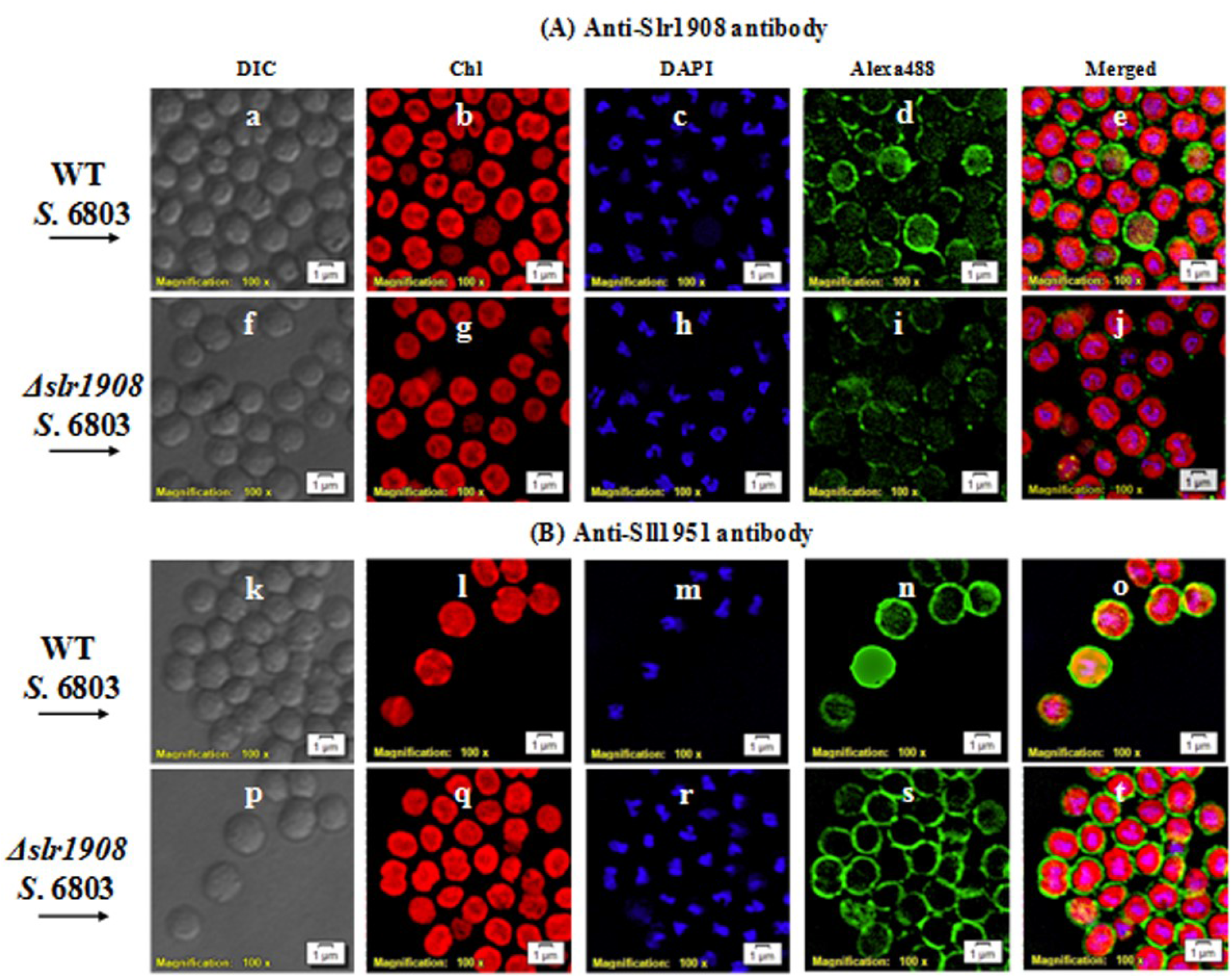
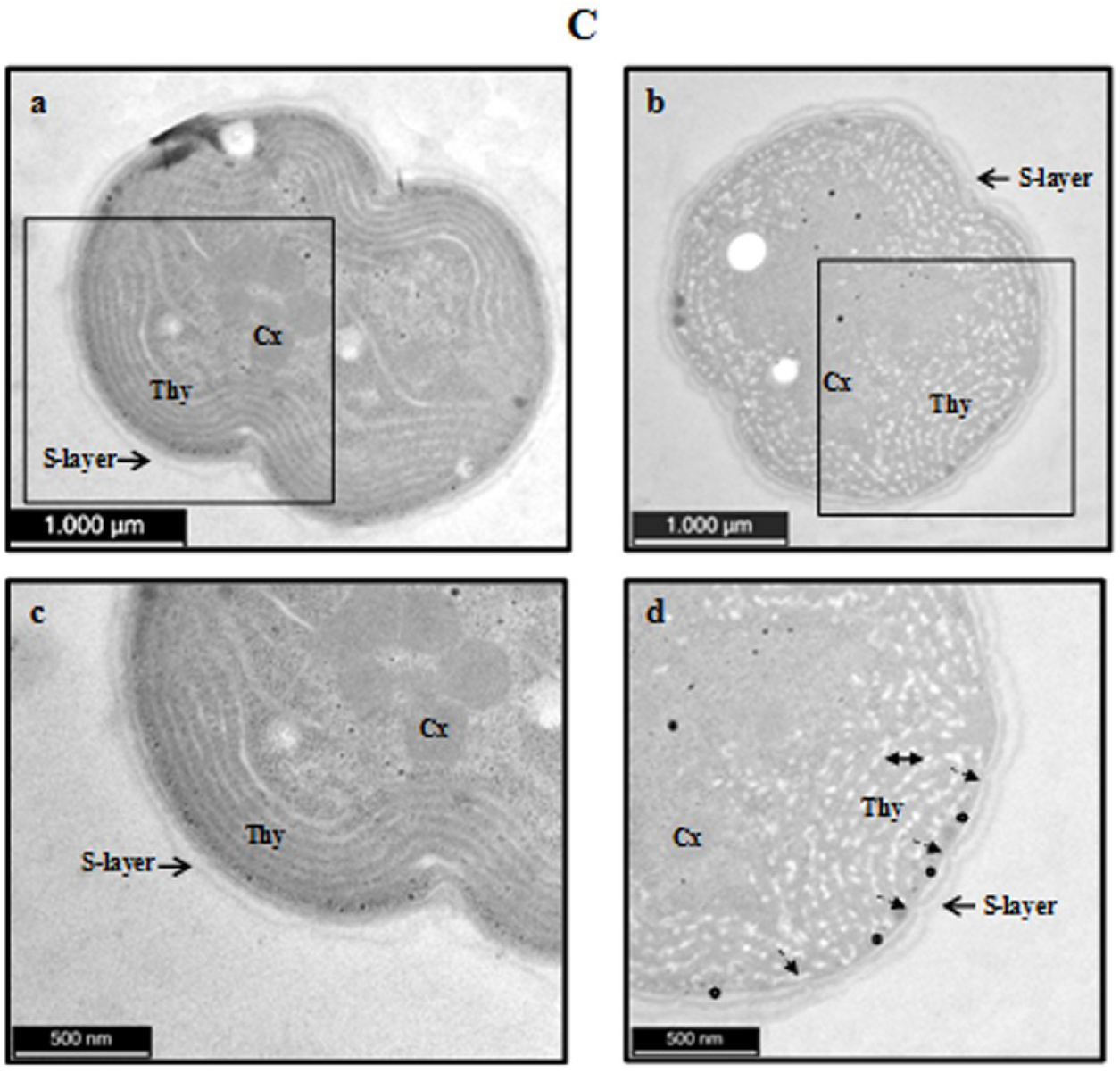
*In vivo* immunolocalisation of **(A)** Slr1908 and **(B)** Sll1951 in WT and *Δslr1908* cells. Immunofluorescence was performed using anti-Slr1908 anti-Sll1951 primary antibody and Alexa-488 conjugated secondary antibody. Cells were located by DIC microscopy (a, f, k, p), phycobilisome/chlorophyll fluorescence (b, g, l, q) and DAPI staining of DNA (c, h, m, r) and Alexa-488 fluorescence (d, i, n, s). 460–495 nm, 530–550 nm and 340–390 nm filters were employed for visualizing Alexa-488, phycobilisome/chlorophyll and DAPI fluorescence, respectively and image was formed by merging red, green and blue channels (e, j, o, t). Scale bar = 1 μm. The experiment was repeated thrice and ∼600 cells were viewed in random fields. Chl: Chorophyll **(C)** Ultrastructure of WT (a and c) and *Δslr1908* cells (b and d). S-layer: Surface layer; Thy:Thylakoids; Cx: Carboxysomes. Wavy cell envelop is marked by black arrow **(↔**). White starch deposits are marked by black double headed arrow ((→). Irregular envelop spacing is shown by a star (⛯). Scale bars measurements are depicted on respective image.

Effect of *slr1908* deletion showed distinct effect on the ultrastructure (Fig. 4 C). The WT cells showed concentric peripheral rings of thylakoids with carboxysomes in the centre and a continuos cell envelop with discernible regularly spaced surface-layer (S-layer) [Fig. 4 C, a and c]. However, the thylakoids were interspersed with abundant white coloured starch deposits in the Δ*slr1908* cells only, indicative of stress [24]. Further, the envelop layers in the Δ*slr1908* cells were wavy with irregular spacing. The ROS content measured as DCHFDA fluorescence (A.U.) on equal cell density basis also substantiated that Δ*slr1908* cells were indeed under higher oxidative stress after one week of growth in comparison to the WT (DCHFDA fluorescence was 4154±330 and 2750±81 A.U. for Δ*slr1908* and WT cells respectively).

### 3.5 One week old Δ*slr1908* cells exhibited ∼80% less Fe, ∼50% less Mn and reduced Fe^3+^ uptake in comparison to the WT

**T**he intracellular concentration of six essential metals (Fe, Mg, Ca, Cu, Mn and Zn) was analysed by atomic absorption spectrophotometry on equal dry weight basis. The nitric acid digest of one week old Δ*slr1908* cells had ∼ 80% less internal Fe in comparison to the WT (Fig. 5A). This deficit was reduced to ∼ 55% in three weeks old Δ*slr1908* cells (Fig. 5A). The intracellular content of Mn was also ∼50% reduced in one week old Δ*slr1908* cells that was restored to WT levels at the end of three weeks (Fig. 5A). Notably, no other tested metal (Cu, Ca, Mg and Zn) showed any significant difference in their cellular quotas between the WT and the Δ*slr1908* cells (Fig. 5A and B).

**Fig. 5:**
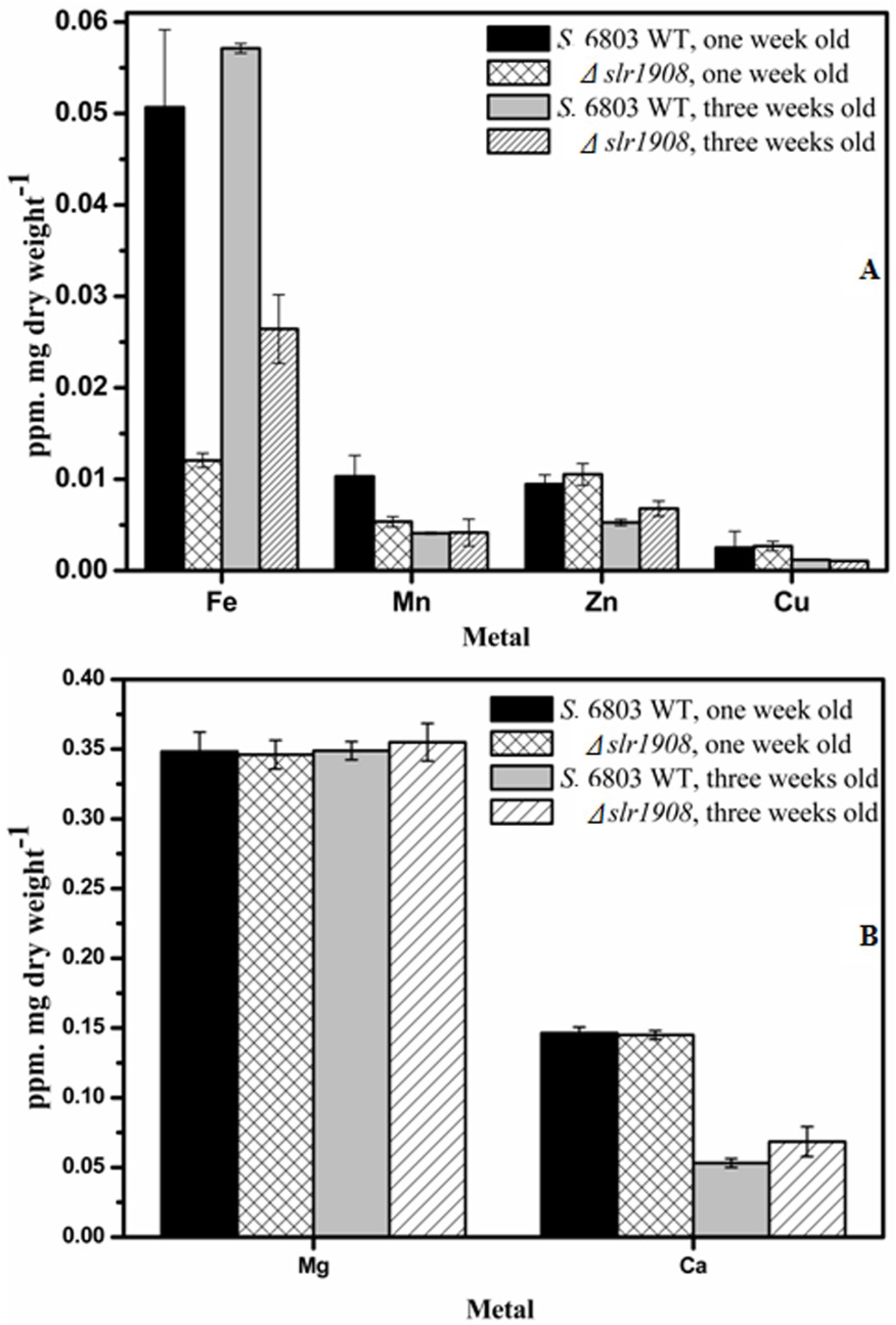
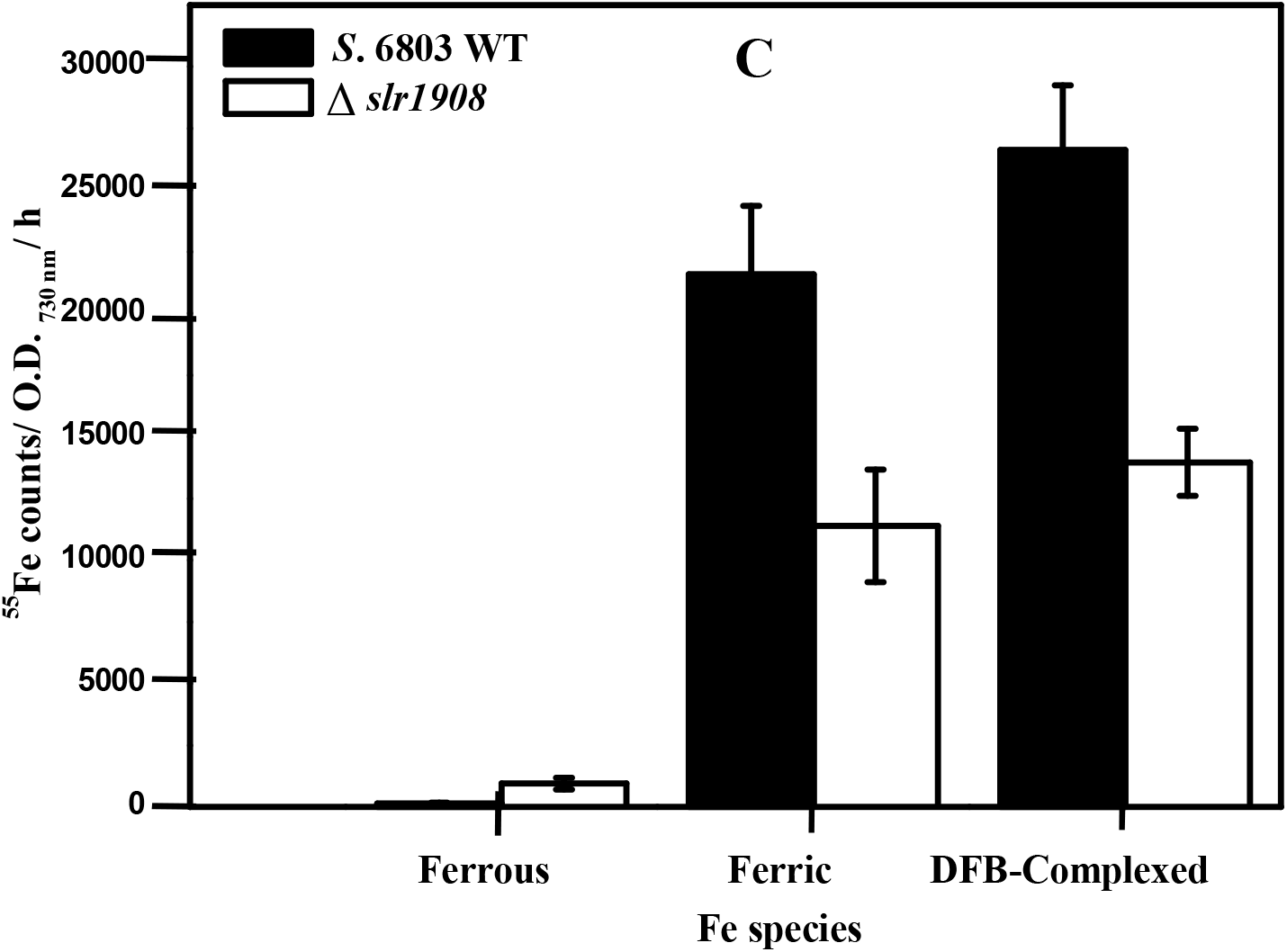
Metal content analysis of WT and *Δslr1908* cells at the end of first week and third week of growth by atomic absorption spectroscopy **(A)** Fe, Mn, Zn and Cu **(B)** Mg and Ca. The experiment was done in technical and biological quadruplets. **(C)** Labelled ^55^Fe uptake by WT and *Δslr1908* cells at the end of first week. The experiment was done in technical and biological quadruplets/ triplicates. Values represent Average±SEM.

Further, in order to elucidate the effect of Fe species on Fe uptake, radioactive tracer ^55^Fe uptake studies were performed on one week old cultures. Fe was kept reduced as Fe^2+^ by adding ascorbate whereas only oxidised form Fe^3+^ remains by adding ferrozine (as ferrozine complexes free Fe^2+^ forming a stable coloured complex). Fe chelated with DFB served as the source of complexed Fe. The results showed that though Fe could be taken up as Fe^3+^ as well as DFB-Fe complex by both cell types, it was ∼ 50% less in the Δ*slr1908* cells in comparison to the WT (Fig. 5C). Fe was poorly taken up by either cell types when supplied as Fe^2+^ and showed a reverse trend wherein Fe^2+^ uptake was seven times more in Δ*slr1908* cells in comparison to the WT cells (Fig. 5C).

### 3.6 Δslr1908 cells showed increase in the amount of ∼ 60 kDa OMP and decrease in content/ activity of Fe requiring proteins

Out of the seven putative OMPs in *S*. 6803 (Table S1), Slr1841 (∼ 67 kDa) and Slr1908 (∼ 64 kDa) are the most abundant at transcript as well as protein level. The reported relative abundance ratio of Sll1951 (S-layer protein), Slr1841, Slr1270 (TolC like OMP), Slr1908, and Slr0042 proteins in purified OM of *S*. 6803 is 1.5:5.5:0.4:2.3:0.3 [17]. Sll1951 being an S-layer protein is co-purified as it interacts with OM by non-covalent interactions and cannot be removed completely unless treated with chaotropic agents [17, 30]. Considering the other four as bonafide OM proteins, the ratio between Slr1908 and Slr1841 turns out to be ∼ 1:2 and that between Slr1908 and Slr0042 ∼ 10:1. In order to ascertain the complete segregation of Δ*slr1908* cells, western blot analysis of the whole cell lysates was performed using anti-Slr1908 antibody (Fig. 6 IA). Anti-Slr1841 and anti-Slr1270 served as positive control [21] (Fig. 6 IB and IC). Bioinformatic sequence alignment had earlier showed that Slr1908, Slr1841, Sll1550 and Slr0042 share ∼ 90% identity in the first 120 amino acids at the N-terminus (Fig. S1A) whereas the 44 amino acid long SLH-domain was > 95% identical (Fig. S1B). As a result, both anti-Slr1908 and anti-Slr1841 antibodies showed cross-reactivity with each another as well as with some lower molecular mass proteins (Fig. 6 IA and IB). The Δ*slr1908* cells did not show any band corresponding to the Slr1908 protein re-iterating complete segregation in both one week and three weeks old cells (Fig. 6 IA). Interestingly, a ∼ 60 kDa protein was prominently induced in the mutant that is most likely to be Slr0042 (as a band lower in size was also observed that can be ascribed to Sll1550, see table S1 for size description of various OMPs) (Fig. 6 IA). Slr1841 and Slr1270 appeared similarly expressed in both the cell types (Fig. 6 IB and IC). The superoxide anion detoxifying Fe-SOD (probed using an anti-Mn-SOD from *Anabaena* 7120, with which it shares ∼ 50% identity) was repressed in the Δ*slr1908* cells after one week and was slightly restored after three weeks on equal protein basis (Fig. 6 ID and IE).

**Fig. 6:**
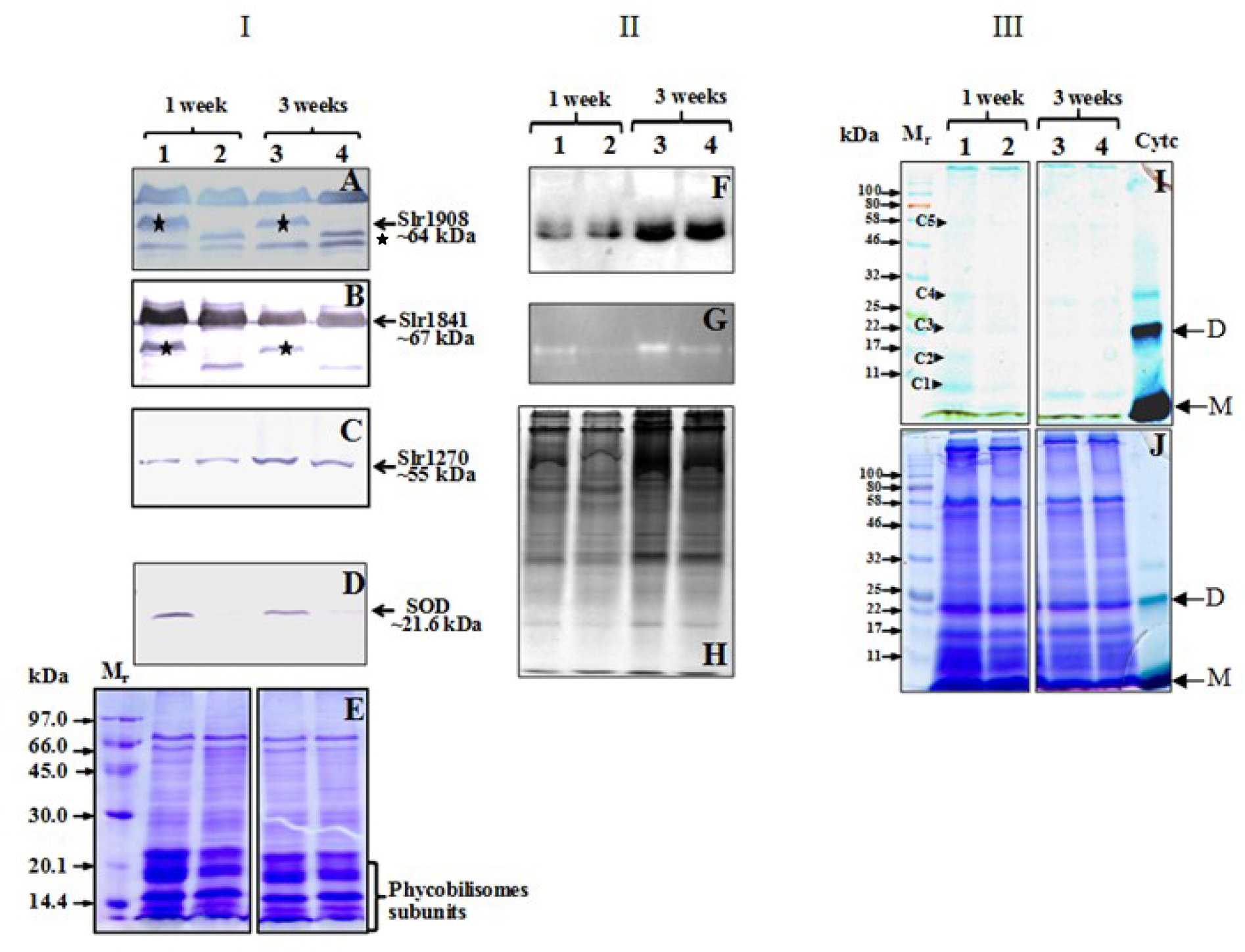
**(I)** Western blot analysis of WT and *Δslr1908* cells at the end of first week and third week of growth (A) anti-Slr1908 antibody (B) anti-Slr1841 antibody (C) anti-Slr1270 antibody (D) anti-Mn-SOD antibody (E) 12% SDS-PAGE showing equal protein loading. The Slr1908 protein is marked by a star (★). The position of the specific band of protein of interest and its observed molecular mass is shown on R.H.S. by a black arrow. **(II)** In-gel activity staining for (F) Catalase (G) SOD and (H) 10% native-PAGE showing equal protein loading. **(III)** In-gel TMBZ staining of whole cell lysates (I) for cytochromes shown as light blue bands. Major cytochrome bands have been marked C1-C5. (J) 12% non-reducing PAGE showing equal protein loading. 10 μg Cytc has been used as a positive control (marked by black arrow on RHS). M: Monomer; D: Dimer. Applies to all panels, Lanes 1 and 3: WT cell lysate at the end of first week and third week of growth; Lanes 2 and 4: *Δslr1908* cell lysate at the end of first week and third week of growth.

As the Δ*slr1908* cells appeared stressed in TEM as well as DCHFDA assay, the activity of ROS detoxifying enzymes catalase and Fe-SOD were also assayed. Although the catalase activity remained comparable in the WT and the Δ*slr1908* cells mutant at both the time points (Fig. 6 IIF), Fe-SOD was severely affected in one week old Δ*slr1908* cells that was partially restored by the end of three weeks (similar to the results obtained in western blot) (Fig. 6 IIG-H). Further total cytochromes determined by TMBZ staining were also lower after one week of growth in the Δ*slr1908* cells in comparison to the WT cells (marked C1-C5, corresponding to Cytb_559_, Cytc_553_, Cytc_550_, Cyt*f* and Cytp450 respectively, Fig. 6 I-J) [31-32]. Similar to SOD activity, the content of cytochromes in the Δ*slr1908* cells became comparable to the WT after three weeks of growth (Fig. 6 III, I-J).

### 3.7 Δ*slr1908* cells showed increased transcription of *slr0042* along with *isiA*-*isiB, sodB* and photosynthetic cytochromes

The aforementioned results indicated that the *Δslr1908* cells are impaired in Fe import and are trying to combat this stress by altering their Fe demand and reducing the synthesis of Fe requiring proteins such as SOD and cytochromes. These results were further strengthened by analysing the transcription status of Fe requiring proteins as well as other porins by RT-qPCR [26]. The C_t_ values were compared with the RPKM values obtained previously for the WT cells by RNA-seq experiment (Table S3). In general, higher the RPKM value, lower was the C_t_ value, indicating experimental reliability. The *isiA*-*isiB* genes were highly transcribed in the Δ*slr1908* cells (∼22 fold and ∼32 fold respectively) in comparison to the WT which is in synchrony with the Δ*slr1908* cells’s Fe deficiency (Fig. 7, see also the inset). Further, out of the four tested OMP genes, *slr0042* was sixteen-fold induced in the Δ*slr1908* cells in comparison to the WT (Fig. 7). Though *slr1841* transcription in the Δ*slr1908* cells also appeared elevated, it did not fall in the 95% confidence interval for the student’s t-test and hence was not considered significant (Fig. 7). The transcription status for *sodB* gene was ten-fold higher in the Δ*slr1908* cells while the *katG* induction was not significantly altered (Fig. 7). Transcription of *bfrA* (*sll1341*) gene required for intracellular Fe storage was not significantly upregulated whereas *mrgA* transcript (*slr1894*), encoding a DPS family protein required for intracellular mobilisation of stored Fe was ∼ 14 fold higher in the Δ*slr1908* cells (Fig. 7) [33-34]. Out of the five tested cytochrome genes, three genes encoding cytochromes of membrane localised photosynthetic complexes i.e., cytc550 (*psbV*, component of PSII), *cytf* (*petA*, component of Cyt*b*_*6*_*f*) and cytb_*559*_ (*psbE*, component of PSII) were ∼ 10 fold upregulated where as cytP450 (*slr0574* encoding a retinoic acid hydroxylase) and *cytc553* (*petJ*, soluble electron carrier between cyt*b*_*6*_*f* and PSI) didn’t show significant change. The global Fe uptake regulator *fur* also did not show any significant transcriptional difference between the WT and the Δ*slr1908* cells (Fig. 7).

**Fig. 7:**
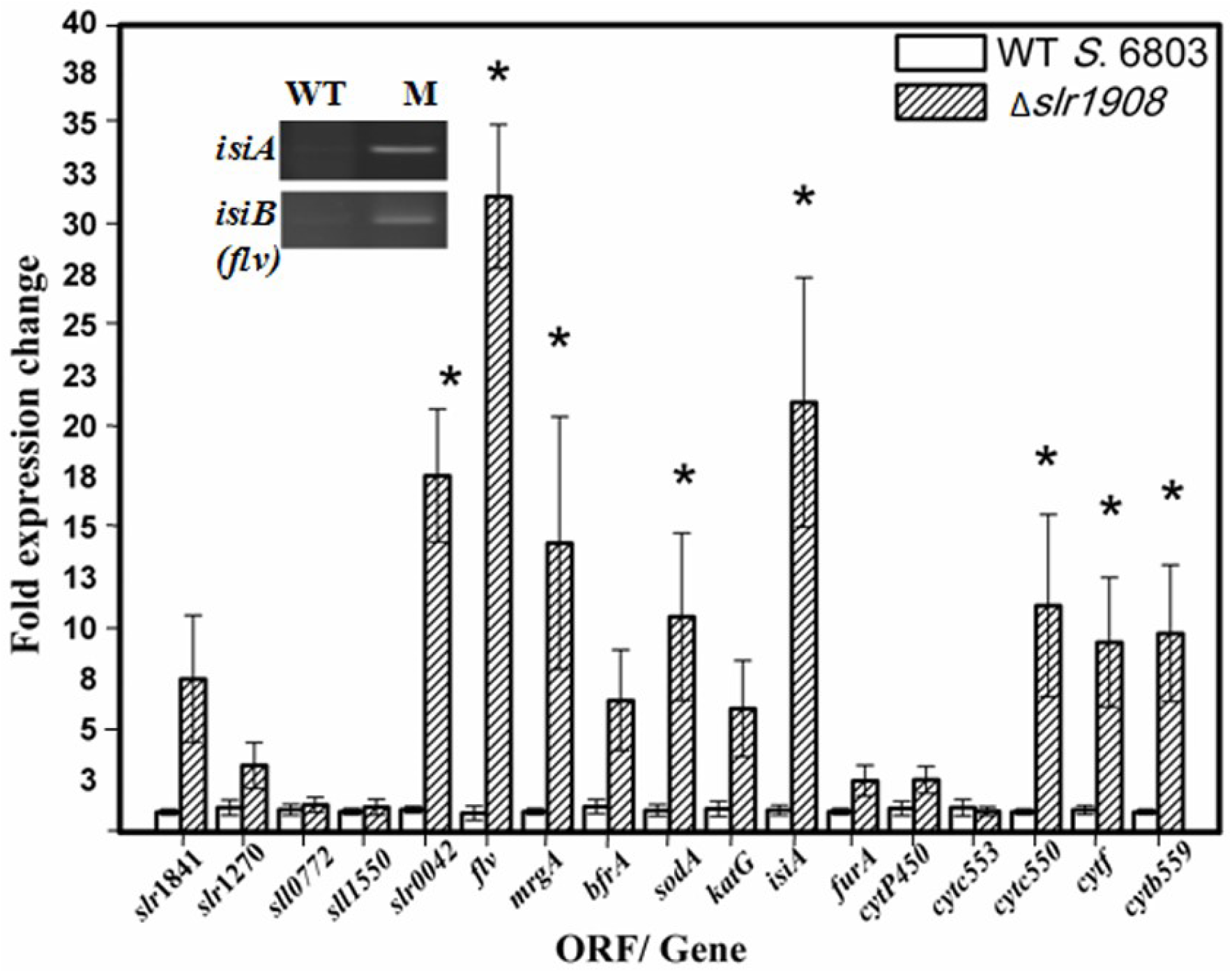
RT-qPCR analysis of expression of OMPs and genes involving Fe in WT and *Δslr1908* cells at the end of first week of growth. The expression analysis was performed according to 2^-ΔΔCt^ method. Values represent average±SEM, where n=5-6. The statistically significant difference as determined by student’s t-test is marked by an asterisk. The inset shows RT-qPCR product of *isiA* and *isiB (flv)* genes resolved on 2% AGE.

## 4. Discussion

Fe represents one of the “necessary evils” of the living world [35]. On the one hand it is indispensable for primary metabolism such as respiration and photosynthesis and on the other hand its high reactivity through Fenton’s chemistry is extremely deleterious to the biomolecules [35]. Indeed, the living world has improvised multiple mechanisms for efficient Fe uptake as well as maintaining its fine intracellular balance [6, 10, 36-38]. Once inside the cell, Fe can be either used for cellular metabolism as co-factor for various enzymes and redox centres or be stored as cellular reserve for future use so as to avoid oxidative damage [33]. Dedicated regulators such as Fur protein orchestrate this fine tuning [39].

There is abundant information on Fe acquisition pathways across bacterial inner membrane [40]. It is also well established that under conditions of Fe deficiency, high affinity TonB-dependent transporters, import Fe primarily as ferric siderophores at the expense of energy through OM [4, 13]. In the case of *E. coli*, there are six reported TBDTs (FhuA, CirA, FecA, PiuA, FepA, FhuE) for the uptake of various Fe-siderophores and their function is believed to be mutually redundant to obtain Fe under a variety of environmental conditions [41]. However even a combined mutant of all six genes showed growth on LB plates and needed 2,2-bipyridine to pose severe Fe limitation and hamper growth, indicating that apart from these TBDTs there may still be other Fe import routes in the organism [41]. Similarly in case of *S*. 6803, four TBDTs and four outer membrane porins have been shown to bring about Fe import across its outer membrane [14]. However, the TBDT quadruple mutants do not appear malnourished when grown under regular BG-11 medium in the laboratory conditions with ∼ 23 µM of Fe indicating that their Fe requirement is already met with by a yet unknown transporter [14]. Further, Fe deficiency symptoms in the mutant as well as upregulation of the corresponding loci in the WT is observed only when the cells are transferred from an Fe sufficient to an Fe limited/ deficient medium [14, 42]. The results suggest that these loci are obligatory to tide over the Fe limitation and there indeed are other proteins involved in Fe homeostasis across the outer membrane. For example, under Fe adequacy in the external environment such as BG-11, Fe may be taken up by specific/ non-specific passive diffusion channels i.e. porins as previously suggested [14]. However, since the quadruple knock out of four poorly expressed porins of *S*. 6803 could also not produce any Fe limited phenotype in standard BG-11 conditions, the search for the elusive OM transporter continued [14]. In the present study, the Δ*slr1908* cells that grows slowly in Fe replete conditions, shows significantly less total intracellular Fe as well reduced Fe uptake and exhibits classical signs of Fe deficiency qualifies to be the “specific” Fe importer in *S*. 6803 in complete BG-11 medium under laboratory conditions, as intracellular concentration of other metals (except Mn) remains the same in the Δ*slr1908* cells with respect to the WT. Since, OM lacks a proton-motive gradient and hence requires energisation through IM potential via TonB for active import, it may be possible that Fe uptake via Slr1908 occurs by diffusion as this protein lacks a TonB box.

Fe has been shown to control intracellular Mn accumulation in *S*. 6803 [38]. Mn and Fe are both transition metals with atomic numbers of 25 and 26 respectively. Though Fe exists in the environment as Fe^3+^ forming insoluble non-bioavailable hydroxides, Mn is predominantly in hydrated Mn^2+^ state which is readily taken up by living organisms [43]. Hence, in spite of being in nano molar range, Mn is not considered a limiting factor in aqueous environment [44]. However, like Fe, OM transport route for Mn are also not known [10]. Each cell of *S*. 6803 an accumulate upto 10^8^ Mn^2+^ in its envelope layer that acts as a pool for maintaining intracellular Mn ions at 10^6^/ cell [45-46]. Under Mn-deficient/ limiting conditions, it is transported through MntABC transporter in the plasma membrane [12, 43]. Due to similar electronic configuration and high spin (d5) states of Mn^2+^ and Fe^3+^, their complexes have similar geometries and hence their promiscuous entry through the same portal (as observed in this study) appears possible and reasonable as the receptor/ transporter might not be able to distinguish between them [38]. Alternatively, the changes is S-layer morphology due to absence of Slr1908, might somehow lead to lower surface bound Mn^2+^ and hence lower intracellular content.

Despite of having 80% less Fe, the mutant cells had twice the oxidative stress in terms of DCHFDA fluorescence that was reinforced in TEM images. This appears contradictory, as Fe supposedly is one major contributory factor for generating oxidative stress through Fenton’s reaction [37]. Further, the mutant also had low ETRII indirectly implying low O_2_ evolution that is another mode of oxidative stress generation. This can be attributed to 50% less Mn concentration in the mutant. Mn is known to protect cells from oxidative stress by several enzymatic as well as non-enzymatic mechanisms [44, 47]. Mn can execute non-enzymatic disproportionation of superoxide in association with small metabolites such as phosphates and carbonates at physiologically relevant concentrations [48]. Interestingly, many enzymatic antioxidants such as SOD and catalase require either Fe or Mn for their activity. Alterations in total intracellular Mn levels can alter ROS sensitivity in bacteria and high intracellular Mn is one of the contributing factors for unusually high oxidative stress tolerance of radioresistant *Deinococcus radiodurans* [47, 49]. Further, the coordinated decrease in Mn quota, can also be the adaptive response to reduce the transport of electrons through PSII.

Besides stress symptoms, the TEM micrographs of the Δ*slr1908* cells showed irregular spacing between peptidoglycan, OM and S-layer which is the outcome of loss of otherwise abundant Slr1908 that has the SLH domain which may be interacting with the peptidoglycan layer in the cell envelop (inherent property of OMP’s SLH domains) thus keeping the cell envelop regularly spaced in the WT. Notably, CRISPR-Cas mediated partial silencing of the other major OMP Slr1841 in *S*. 6803 (having SLH domain) generates an OM-deprived phenotype indicating the importance of these proteins in maintaining the cell envelop integrity [50].

The Δ*slr1908* cells though initially defective in Fe (and Mn) uptake, have been modulating its intracellular machinery to abrogate the stress and enhance Fe (and Mn) import. Out of the four OMPs, the Δ*slr1908* cells had an upregulation of porin Slr0042 at both RNA and protein level. A role of Slr0042 in Fe acquisition in the quadrapule OMP mutant has been established previously in *S*. 6803 and hence its up-regulation seems logical to take up more Fe [14]. However, this OMP is not as efficient as Slr1908, as Fe deficiency symptoms persisted even at the end of three weeks. Also, Mn content in Δ*slr1908* cells became comparable to the WT at the end of three weeks that is exhibited also in its enhanced growth rate in the later half. This better growth may be ascribed to increased Mn and/ or Fe content that may be resulting from Slr0042 induction or up-regulation of yet other OMPs such as TBDTs, that needs to be explored further.

Of the two Fe requiring oxidative stress detoxifying enzymes, SOD and catalase, the mutant is mostly allocating the required Fe pool to catalase so as to detoxify the long lived H_2_O_2_ that may form hydroxyl radical whereas the superoxide dismutase, though transcriptionally enhanced, remained poorly translated and hence exhibited less content as well as activity in the mutant. Due to short lived nature of superoxide anion, it is unlikely to cause large destruction to the cellular machinery by itself. However, the long lived H_2_O_2_ that can undergo Fenton’s reaction with Fe itself thus producing highly reactive hydroxyl free radical might prove more destructive. This modulation of proteome might be a part of “Fe sparing response” in which the translation of proteins that require Fe in large quantity are repressed/ blocked in order to channel the limited intracellular Fe towards the most essential Fe utilising proteins [51]. This is also supported by comparing photosynthetic cytochromes *cytc*_550_, *cytf* and *cytb*_*559*_, which though transcriptionally upregulated are not observed in higher amounts at protein level in TMBZ staining. They are required for photosynthesis and hence are essential as the organism is maintained only under photosynthetic conditions [14]. However, their translational down-regulation seems mandatory for fair distribution of available Fe for simultaneous balance between maintaining metabolism and abrogating stress.

It has been previously shown that proteoliposomes reconstituted with OMPs of WT and *Δslr1270* (containing mainly Slr1841, Slr1908 and Sl0042) cells are capable of permeating only small inorganic molecules such as sodium nitrate and disodium phosphate and further experimentation confirmed that Slr1270 is responsible for moderate permeability of organic solutes upto a size of ∼ 666 Da [14, 17]. However, such in vitro experiments cannot really mimic the actual *in vivo* molecular conformation as well as the change that it may acquire while interacting with the substrate as well as negotiating with other participating neighbouring lipids and proteins. It is interesting to note that Fe in the BG-11 medium is supplied as ferric ammonium citrate (FAC) with a molecular mass of 262 Da. Additionally EDTA is added in the medium that will chelate any available free Fe yielding a complex of molar mass of ∼ 367 Da. Thus, the FAC and Fe-EDTA might be able to enter through a general diffusion porin which have a size cut off of ∼ 600 Da, beyond which the molecules are not able to enter the cell. Considering that *in vitro* permeation experiments showed poor permeability of Slr1908, Slr1841 and Slr0042 to small organic substrates such as glucose (Molar mass of 180 Da) whereas the *Δslr1908* mutant in this study seems to be defective in taking up complexed Fe (as FAC in the BG-11 medium) we hypothesize that the protein might be changing its pore size *in vivo* in response to extracellular complexed Fe. OMPs are known to have a plug domain which can alter its position for modulating the pore size in response to export/ import signal.

High transcription of *slr1908* under laboratory conditions in *S*. 6803 [16, 27] clearly signifies its important biological function such as Fe import as shown in this study. *S*. 6803 is a fresh water cyanobacterium and unlike its open ocean relative such as *Synechococcus*, it has genes for both TBDTs and porins for Fe uptake [14]. These multiple modes of Fe uptake are a suitable adaptive strategy for efficient Fe uptake under varying Fe regimes i.e., when Fe is readily available, the cells may only need to rely on porins for passive Fe uptake. Though Fe bioavailability is a severe growth limiting factor for the organisms, there are reports of Fe content as high as 4-30 μM along with high organic matter content in surface water of various fresh water lakes [52]. In such Fe rich habitats, it would be beneficial for the organism to take up Fe via a passive mechanism such as Slr1908 porin.

## Acknowledgement

Authors would like to thank Analytical Chemistry Division, BARC for atomic absorption spectroscopy, Dr. Prashanth Raghavan, Molecular Biology Division, BARC for providing Mn-SOD antibody, and Dr. Amit Prakash Shrivastav, Material Science Division, BARC in acquiring TEM micrographs.

## Author Contributions

**RA:** Conceived and designed the project, executed the experiments, analysed the data and wrote the MS, **SDC:** Helped in low temperature fluorescence experiments, **AVSSNR:** Analysed the data and reviewed the MS.

## Abbreviations

AAS: Atomic absorption spectroscopy
DCHFDA: 2′,7′-Dichlorofluorescin diacetate
ETR: Electron transport rate
FAC: Ferric ammonium citrate
GT: Glucose tolerant
NBT: Nitro blue tetrazolium
OMP: Outer membrane protein
PAM: Pulse amplitude modulated
PPFD: Photosynthetic photon flux density
RT: Room temperature
RT-qPCR: Reverse transcriptase quantitative polymerase chain reaction
SLH: S-layer homology
SOD: Superoxide dismutase
TBDT: TonB dependent transporter
TEM: Transmission electron microscopy
TEMED: Tetramethylethylenediamine
TMBZ: 3,3’,5,5’-Tetramethylbenzidine
TU: Transcriptional unit
UTR: Untranslated region
WT: Wild type

