## Supplementary data for "Iron acquisition across outer membrane: The role of major outer membrane protein Slr1908 under iron supplemented conditions in *Synechocystis* 6803"

Running title: *in vivo* role of *Slr1908* in *Synechocystis* 6803

| **Table S1: List of putative OMPs in the genome of *Synechocystis* PCC 6803 (from: http://genome.microbedb.jp/CyanoBase)** | | | | | | | |
| --- | --- | --- | --- | --- | --- | --- | --- |
| **ORF** | **Chromosomal location** | **No. of bases** | **No. of amino acids** | **Signal peptide**  **(SP)** | **Size after SP removal**  **(kDa)** | **pI after SP removal** | **SLH domain** |
| *slr1272* | 1864239….1865003 | 765 | 254 | 1-28 | 25.246 | 4.66 | 70-114 |
| *sll1550* | 3454422….3456056 | 1635 | 544 | 1-26 | 55.992 | 4.41 | 61-105 |
| *sll0772* | 2397478….2399118 | 1641 | 546 | 1-21 | 57.146 | 4.9 | NP |
| *slr0042* | 3171428….3173158 | 1731 | 576 | 1-27 | 58.958 | 4.46 | 61-105 |
| *slr1908* | 611057….612832 | 1776 | 591 | 1-28 | 61.634 | 4.9 | 80-124 |
| *sll1271* | 1112531….1114249 | 1719 | 572 | 1-28 | 60.101 | 4.48 | 83-120 |
| *slr1841* | 958137….960029 | 1893 | 630 | 1-27 | 64.993 | 4.4 | 61-105 |
| **Actual ORF no., its chromosomal location, gene size and no. of amino acids in full length protein have been given. Further position of signal peptide (SP), size after SP removal (in kDa, isoelectric point (pI) and position of S-layer homology (SLH- domain) have been listed based on prediction by http://smart.embl-heidelberg.de/. NP: not predicted** | | | | | | | |

| **S. No.** | **Table S2: List of primers used in this study** | |
| --- | --- | --- |
|  | **Primer name** | **Primer sequence (5’ 3’ )**  **(Restriction sites in bold italics)** |
| 1. | 1908_F (P1) | TCGA ***CATATG*** AATAAGTTGACCAGTCATTTAC |
| 2. | 1908_R (P2) | TCGA ***CTCGAG*** GAACTTGAAGGTGGTGC |
| 3. | 1908_flanking_F (P3) | GAATTTCTTGGTCTTCGCTGAT |
| 4. | 1908_flanking_R (P4) | CGACGAACTGTACCGTCTAAAT |
| 5. | 1908_RT_R (P5) | GGTGGTGGTGGAGAATTGAT |
| 6. | rnpB_136_F (P6) | CGTTAGGATAGTGCCACAG |
| 7. | rnpB_136_R (P7) | CGCTCTTACCGCACCTTTG |
| 8. | 1907_200_F (P8) | GAAGCCCTACCATGCTGAAT |
| 9. | 1907_200_R (P9) | CCATGGCGATGGGATTATCA |
| 10. | 1909_200_F (P10) | CAACGTGGGAGCGGATATTT |
| 11. | 1909_200_R (P11) | CCTGGGCTACCAATTGAATCAC |
| 12. | SOD_200_F | GACGTAATTAAAGCCGTTGCTG |
| 13. | SOD_200_R | GCTCCAGCTTGTTTGAAAGC |
| 14. | Kat_200_F | ACCACGAATGGGAACTGAAG |
| 15. | Kat_200_R | ATAATCCGGCTCCCGATAGA |
| 16. | isiA_200_F | GTGGATACCTATCCCTACTTTGTC |
| 17. | isiA_200_R | CCCAGGAACAGGAGATGATG |
| 18. | Flv_200_F | TGCCTGATTATCGGTTGTCC |
| 19. | Flv_200_R | TTACCGCCTAATCCACTGATTT |
| 20. | C553_200_F | CTGGGTAACACCGCCCTA |
| 21. | C553_200_R | CCAGGCATAGCGCCATTA |
| 22. | C550_200_F | GACACCTGCACCCAATGT |
| 23. | C550_200_R | TAGATGTCGGGACGGGAAATA |
| 24. | P450_200_F | AAGGGCTATTTAGTCTGCCAAT |
| 25. | P450_200_R | CTTCAGCTCCGGCAAAGATA |
| 26. | Cytf_200_F | ACCTACTTCCAGCCCTACC |
| 27. | Cytf_200_R | CTCAGCAAACCAGTGGGATAA |
| 28. | B559_200_F | TTCCGATATTGTCACCAGCAT |
| 29. | B559_200_R | GAATTTCCTGATTAATGTCGTAGCG |
| 30. | furA_200_F | TCAGTGCCGAGGAGTTACA |
| 31. | furA_200_R | GCGATTACATTGCACACAGAC |
| 32. | mrgA_200_F | AGCCGATACCTACACCCTTTAT |
| 33. | mrgA_200_R | CCGAACTCAGTTTAGCGAACTC |
| 34. | bfrA_200_F | ATGCCAGCCTGTTGATTGA |
| 35. | bfrA_200_R | CAATAGGTCCCGGCTTTGATAG |
| 36. | 1841_200_F | GTGGTGGTTCTGTTACTAACCC |
| 37. | 1841_200_R | ACCTTGAGTGCCACCTTTAC |
| 38. | 1270_200_F | GAGCAACAGGAAGCTGGTTTA |
| 39. | 1270_200_R | GTCAGGTTGATGTTACCCACTG |
| 40. | 0772_200_F | CAACTGAGTGTGGGCTACTT |
| 41. | 0772_200_R | TGAATGGCCAACAGACTACC |
| 42. | 1550_200_F | TACTTTCCAGCCAGTGTTAGG |
| 43. | 1550_200_R | GGGTGCCAAAGGTAGTCAAT |
| 44. | 0042_200_F | GCTGGTCTCAATGCCTGTAT |
| 45. | 0042_200_R | AGTTTGGTGGTGGTGGAAA |

Table S2: List of primers used in this study for PCR/ RT-qPCR amplification of the corresponding locus as defined in the primer name. The italicized sequences (wherever applicable) denote the restriction sites incorporated in the primers to assist in cloning. F: Forward primer; R: Reverse primer. Primer names in parentheses (P1-P11) are used in Fig. 1 and also elaborated in the text, wherever applicable.

| Table S3: **R**eads **P**er **K**ilobase of transcript per **M**illion mapped reads (RPKM) values of WT (from RNA-seq experiment) and C_t_ values of WT and *Δslr1908* cells (from RT-qPCR experiment) | | | | | |
| --- | --- | --- | --- | --- | --- |
| ORF | Gene | RPKM values (WT)  (Mean±SEM) | WT-C_t_  (Cycle number)  (Mean±SEM) | *Δslr1908*-C_t_  (Cycle number)  (Mean±SEM) | M_r_  (Molecular mass, kDa) |
| *NA* | *rnpB* | NA | 12.0±0.3 | 10.6±0.2 |  |
| *slr1841* | OMP | 10800.6±893.9 | 22.6±0.2 | 18.7±0.7 |  |
| *slr1270* | OMP | 29.5±7.4 | 27.8±0.5 | 25.1±0.7 |  |
| *sll0772* | OMP | 32.6±6.7 | 27.8±0.2 | 26.1±0.6 |  |
| *sll1550* | OMP | 31.8±3.5 | 31.8±0.6 | 29.2±0.5 |  |
| *slr0042* | OMP | 110.6±43.4 | 26.7±0.3 | 22.1±0.7 |  |
| *sll0248* | *isiB* | 23.5±6.8 | 30.4±0.6 | 22.7±0.7 |  |
| *sll0247* | *isiA* | 1347.6±133.8 | 26.2±0.2 | 21.5±0.7 |  |
| *slr1894* | *mrg* | 186.8±10.8 | 24.7±0.7 | 21.2±0.8 |  |
| *sll1341* | *bfr* | 95.2±14.5 | 23.7±0.2 | 19.4±0.8 |  |
| *slr1516* | *sodB* | 4813.6±434.6 | 26.3±0.3 | 22.9±0.8 |  |
| *sll1987* | *katG* | 56.0±4.9 | 27.2±0.4 | 21.5±0.5 |  |
| *sll0567* | *fur* | 61.8±8.9 | 26.9±0.1 | 24.3±0.6 |  |
| *slr0574* | *cyp* | 23.8±3.7 | 28.7±0.4 | 26.0±0.5 | *50.578* |
| *sll1796* | *petJ* | 235.8±71.7 | 27.4±0.3 | 25.9±0.4 | *12.463* |
| *sll0258* | *psbV* | 2099.8±215.0 | 24.5±0.3 | 20.3±0.8 | *15.119* |
| *sll1317* | *petA* | 170.0±36.9 | 27.6±0.5 | 23.2±0.6 | *35.23* |
| *ssr3451* | *psbE* | 591.5±163.1 | 25.0±0.3 | 20.6±0.6 | *9.4* |

**Fig. S1A**

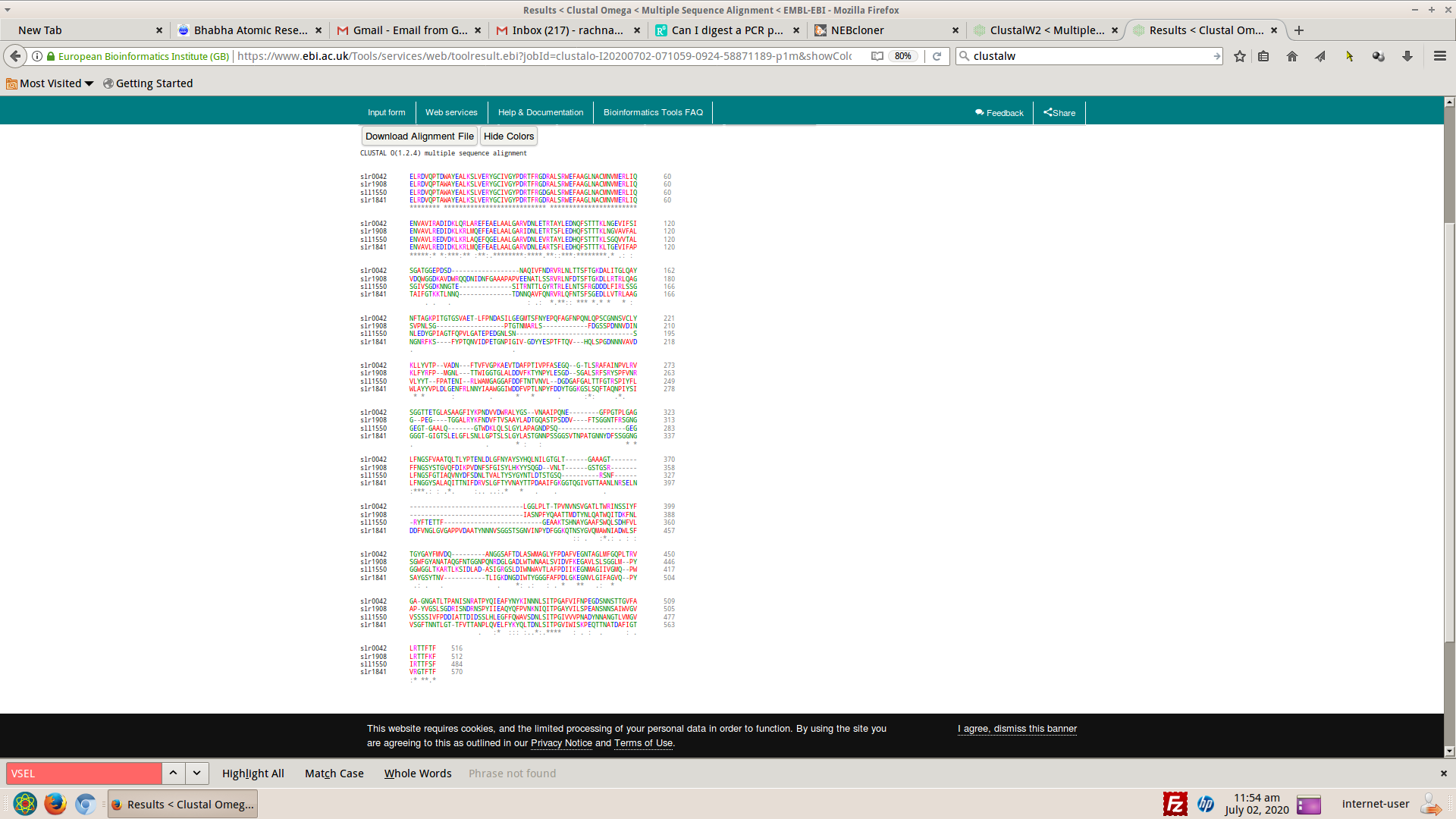

**Fig. S1B**

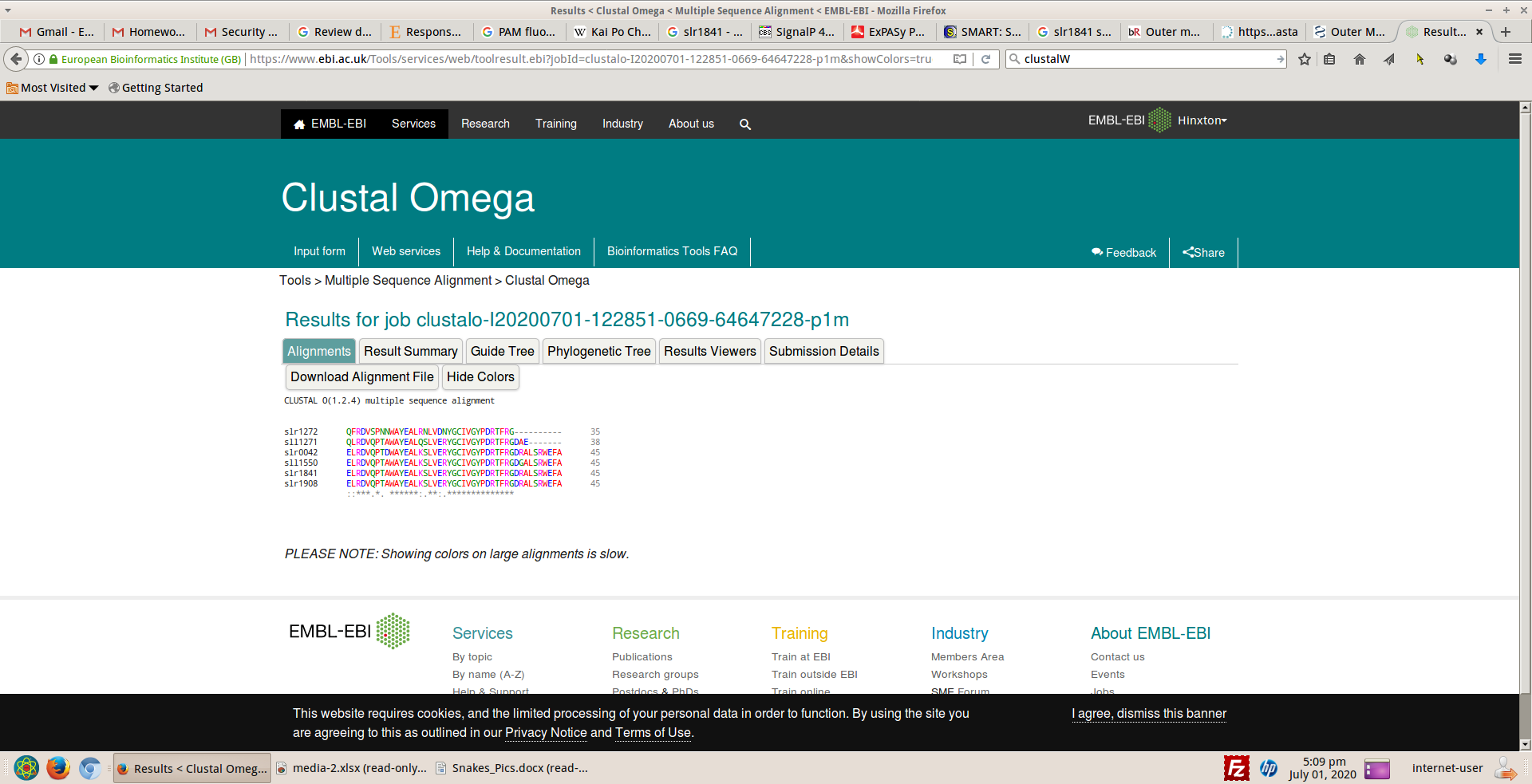

Fig. S1: Multiple sequence alignment performed at https://www.ebi.ac.uk/Tools/msa/clustalo/ **(A)** Full length mature OMPs (after cleavage of signal peptide) showing ~ 80-90% identity between first 120 amino acids at N-terminus **(B)** S-layer homology (SLH) domains of four OMPs showing ~90-95% identity.

**Fig. S2**

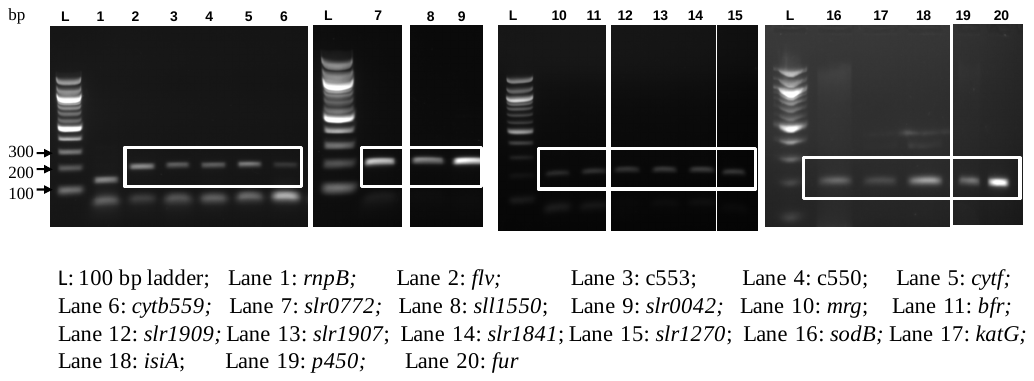

Fig. S2: PCR confirmation of the specificity of the primers for further use in RT-qPCR. PCR was set up using Taq-DNA polymerase, genomic DNA template and primers listed in table S2 and the product was resolved on 2% AGE. The specific product is boxed white. The lane annotation is provided below the gel. bp: base pair
